# Contrast invariant tuning in human perception of image content

**DOI:** 10.1101/711804

**Authors:** Ingo Fruend, Jaykishan Patel, Elee D. Stalker

## Abstract

Higher levels of visual processing are progressively more invariant to low-level visual factors such as contrast. Although this invariance trend has been well documented for simple stimuli like gratings and lines, it is difficult to characterize such invariances in images with naturalistic complexity. Here, we use a generative image model based on a hierarchy of learned visual features—a Generative Adversarial Network—to constrain image manipulations to remain within the vicinity of the manifold of natural images. This allows us to quantitatively characterize visual discrimination behaviour for naturalistically complex, non-linear image manipulations. We find that human tuning to such manipulations has a factorial structure. The first factor governs image contrast with discrimination thresholds following a power law with an exponent between 0.5 and 0.6, similar to contrast discrimination performance for simpler stimuli. A second factor governs image content with approximately constant discrimination thresholds throughout the range of images studied. These results support the idea that human perception factors out image contrast relatively early on, allowing later stages of processing to extract higher level image features in a stable and robust way.

---

Processing in higher levels of the visual system becomes successively less sensitive to image features such as contrast [1, 2, 3, 4] or rotation [5]. In particular higher levels of visual processing seem invariant to a range of image transformations, including contrast changes, rotation, scaling, or different illumination [6, 7, 8].

In recent years, machine learning methods known as deep artificial neural networks [9] have achieved impressive performance on benchmark tasks that require invariant object recognition [10, 11, 12]. Some authors have argued that these algorithms not only achieve impressive performance on invariant object recognition, but also do so in ways that resemble mechanisms of human perception [13] or at least mechanisms in the human visual system [14, 15]. However, more recent findings suggest that processing in deep artificial neural networks differs from processing in the human visual system in significant ways. Specifically, human object recognition seems to largely rely on object shape, while deep networks mostly rely on local texture information [16, 17, 18].

Here, we use deep artificial neural networks in a different way. Instead of directly comparing processing in a deep artificial neural network to processing in the human visual system, we use a deep artificial neural network to *generate* naturalistic-looking images. Specifically, we use a generative adversarial network (GAN, [19]) to approximately constrain high-dimensional stimulus manipulations to the domain of natural images [20].

GANs have recently been successful at generating extremely realistic looking images (see for example, [21, 22]). Furthermore, GANs seem to recover a parametrization of the manifold of natural images that matches aspects of human perception [20] and predicts neural responses in non-human primates [23]. By expressing manipulations of naturalistic images along this approximation of the manifold of natural images, we can characterize complex invariances in human visual perception without a need to rely on analogies between human processing and processing in deep artificial neural network algorithms.

## Results

### Generative adversarial models recover polar-like embedding for natural images

We trained a Wasserstein-GAN [24] with gradient penalty [25] on the 60,000 images of the CIFAR10 dataset [26]. See Methods for details. In brief, a GAN is trained by a game theoretic approach (see Figure 1a), where a generator network *G* maps random vectors from a 128-dimensional latent space to image space and a discriminator network *D* attempts to tell these generated fake images apart from real images. In the following, we refer to the space of 128-dimensional random vectors as the “latent space” of the GAN.

**Figure 1:**
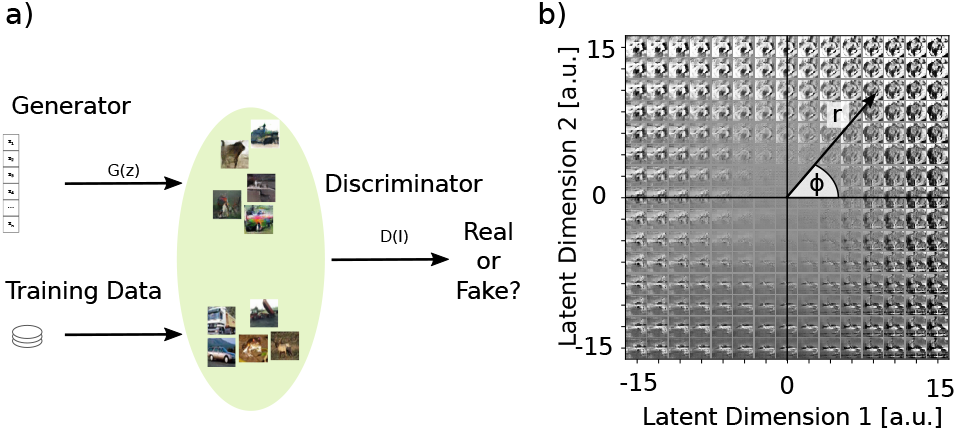
Training generative adversarial networks recovers a latent space of images. (a) Generative adversarial nets are trained using a game theoretic approach. The generator network maps latent vectors 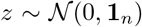 to fake images 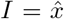 and the discriminator network attempts to tell these images apart from real images *I* = *x*. In the GAN framework, the generator is trained to increase the classification error of the discriminator and the discriminator is trained to decrease its classification error. (b) Illustration of latent space representation recovered by a GAN. The two axes are arbitrary directions in the GAN’s latent space. Each image corresponds approximately to an image drawn from that location in latent space. Superimposed are the variables manipulated in the following studies. The radius *r* is the distance from the origin of the latent space defined by the GAN, the rotation angle *ϕ* describes an *n*-dimensional rotation in the latent space.

Figure 1b shows images generated by passing latent space vectors through the generator of the GAN. We observe that these images had a structure that was well-represented in a polar coordinate system. close to the origin of the GAN’s latent space, images are low contrast, and contrast increases as distance away from the origin inreases. Importantly, images along rays from the origin seem to only differ in contrast, but have the same content. On the other hand, rotations around the origin result in changes in image content, while contrast remains approximately constant.

Yet, by simply looking at Figure 1b, it is difficult to judge if previous notions of contrast are consistent with the length of the the corresponding latent vector’s radius *r*, based on a qualitative impression alone. Furthermore, Figure 1b visualizes a 2 dimensional cut through the 128-dimensional latent space of the GAN. Rotations *ϕ* around the origin would generally be performed in 128 dimensions (where they correspond to 128×128 orthonormal matrices). We therefore performed psychophysical experiments to further characterize the relationship between human perception and the latent space recovered by a GAN.

### Sensitivity to latent space radius follows power law

When observers are asked to discriminate between a stimulus of (superthreshold) contrast *C* and another stimulus of contrast (*C* +Δ*C*), the contrast increment required for threshold performance is proportional to a power *C*^*γ*^ of the pedestal contrast *C* [27, 28, 4]. We reasoned that if latent space radius *r* reflects a stimulus property that resembles image contrast, increment thresholds for radius discrimination should obey a power law as well.

To test this hypothesis, we randomly sampled stimuli from a 128-dimensional hypersphere of radius *r* and then constructed corresponding stimuli by scaling up the latent space representation by (*r* + Δ*r*)/*r*, while keeping the angle of the latent space vector the same (see Figure 2a). This resulted in two latent vectors per trial with radius *r* and *r* + Δ*r* respectively. Observers were asked to detect the image with higher contrast in a spatial 2 alternatives forced choice task (Figure 2b). For fixed *r*, we then varied the increment Δ*r* and found that response accuracy increased with Δ*r*, tracing out a psychometric function (see Figure 2c).

**Figure 2:**
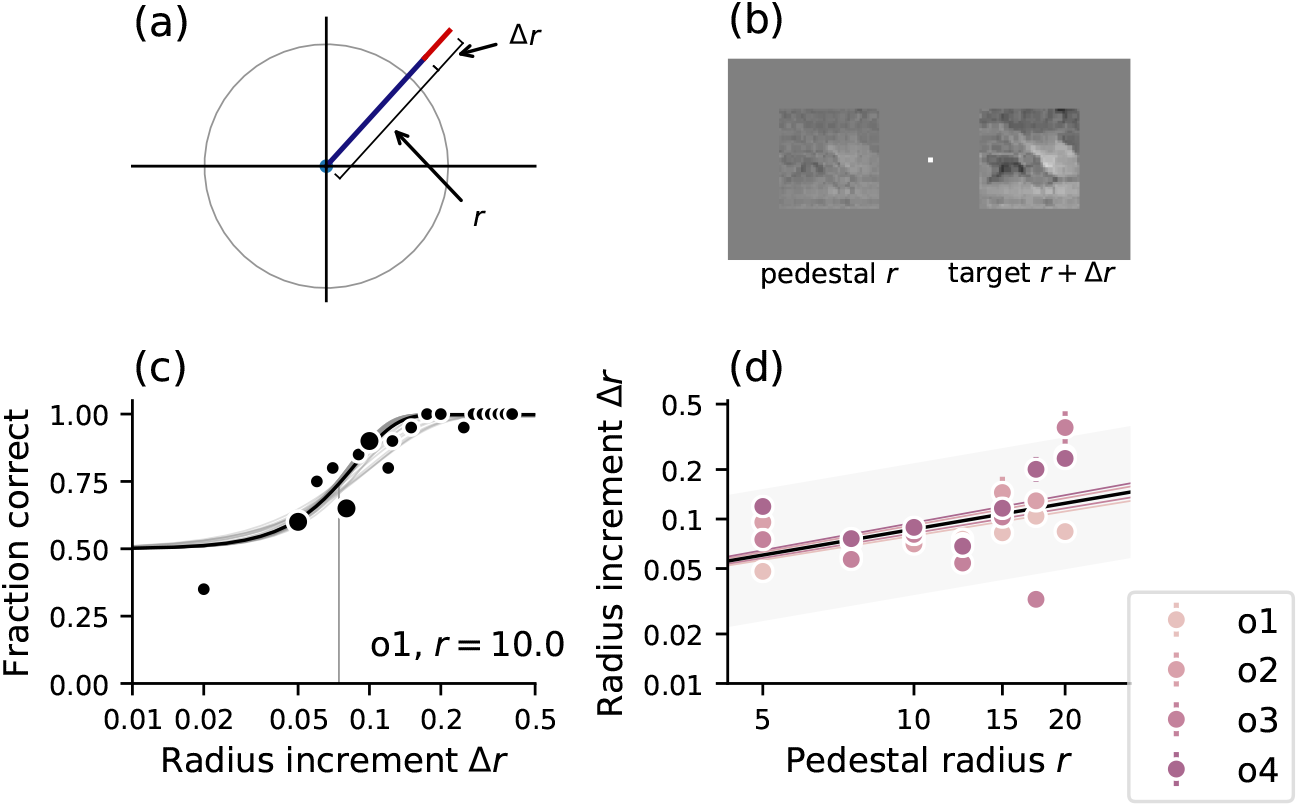
Latent space radius follows a power law. (a) On every trial, the observer sees two images that correspond to the same latent space direction but with different radii. The circle indicates a latent space hypersphere of constant radius, projected to a two-dimensional subspace. A pedestal vector of length *r* is marked in blue and a target increment Δ*r* in red. The target image corresponded to a vector of length *r* +Δ*r*. Latent space directions in different trials were chosen randomly. (b) Example experimental display. Latent space vectors were projected to image space and observers had to choose the image with higher contrast (on the right in this case), while fixating a small dot at the center of the screen. The text below the gray display was not shown during the experiment. (c) Example psychometric function for observer o1 at pedestal radius *r* = 10. Small dots correspond to blocks of 20 trials, larger dots correspond to blocks of 40 trials. The black line is the best fitting psychometric function and light gray lines are samples from the posterior distribution over psychometric functions to illustrate uncertainty in psychometric function estimate. The vertical gray line marks the threshold for this observer. (d) Estimated increment thresholds as a function of pedestal radius. Each dot corresponds to a single threshold (errorbars indicate ± posterior standard deviation), lightly colored lines correspond to best fitting power law functions for individual observers, where different colors indicate different observers. The black line marks the grand average power law function across all observers and the gray area in the background marks a 95% prediction interval for the grand average.

In general, the threshold increment increased as the pedestal radius *r* was increased (Spearman *ρ* = 0.53, *p* < 0.002 permutation test, see dots in Figure 2d). In order to quantify the relationship between pedestal radius and threshold increment, we fit a linear mixed-effects model to the data from all observers (see Methods for details). We found that, on average across observers, increment thresholds increased by 0.523±0.191 log units (mean ± standard error) when the pedestal radius increased by 1 log unit (Wald-test *z* = 2.74, *p* < 0.006), consistent with a power law with exponent 0.523 ± 0.191. Observer specific components for these slopes were negligible (largest slope difference to mean was for observer o1 with −0.033 ± 0.038).

### Latent space radius predicts performance better than other measures of contrast

The results from the previous section suggest that an image’s radius in the latent space of a GAN is closely related to image contrast. To validate that the latent space radius really captured the full contrast discrimination behaviour in our experiment, we investigated whether other measures of image contrast would provide better explanations of behaviour in our task. To do so, we compared five different logistic regression models [29] for the observers’ trial-by-trial response behaviour. Specifically, we compared models that predicted responses from differences in the latent space radii, and from differences between the two image alternatives in Michelson contrast or RMS contrast or Weber contrast. A fifth model predicted responses from differences in all four of these factors; latent space radius, as well as Michelson, RMS and Weber contrast. To prevent overfitting, we regularized the squared norm of the model coefficients (ridge penalty). We tried different weightings of likelihood and penalty, and Table 1 shows the best fitting results for each of the four candidate models based on 5-fold cross validation.

**Table 1:** Consistency between human responses and trial-by-trial predictions based on different contrast measures.

| Model features | o1 | o2 | o3 | o4 | Average |
| --- | --- | --- | --- | --- | --- |
| Radius | 0.81±0.048 | 0.79±0.042 | 0.82±0.031 | 0.83±0.038 | 0.81±0.0087 |
| Michelson | 0.50±0.004 | 0.50±0.002 | 0.52±0.004 | 0.60±0.001 | 0.53±0.0209 |
| RMS | 0.54±0.042 | 0.51±0.020 | 0.53±0.020 | 0.60±0.002 | 0.55±0.0167 |
| Weber | 0.50±0.019 | 0.51±0.007 | 0.52±0.002 | 0.60±0.004 | 0.53±0.0200 |
| All | 0.81±0.049 | 0.79±0.049 | 0.83±0.033 | 0.83±0.039 | 0.81±0.0084 |

Individual observer consistencies are mean and standard deviation across 5 cross validation folds. Average consistencies are mean and S.E.M. across observers.

Across all observers, the model based on GAN latent space radius was consistent with human trial-by-trial responses in about 81% of the cases. Predictions from other contrast measures were generally much worse, with prediction accuracy only marginally above chance (average 53-55%, S.E.M.≈2%). A model that had access to all the latent space radii as well as all three contrast measures did not achieve better prediction performance than a model that predicted trial-by-trial responses from only the GAN’s latent space radius. Other contrast measures therefore only provided negligible information beyond the GAN’s latent space radius. This confirms that a GAN’s latent space radius is indeed a meaningful measure of image contrast for the naturalistic images used in this experiment.

### Constant sensitivity to latent space rotations

In addition to the radius of a point in the GAN’s latent space, we investigated how sensitive observers were to rotations of latent space vectors around the origin. Figure 1 suggests that a GAN’s latent space might have a polar organization, where radius is related to contrast and rotations around the origin correspond to changes in an image’s content. If a GAN factors out contrast from the perceptual representation of the generated images, we would expect that discrimination of images that correspond to slightly rotated latent vectors should not depend on the radius of those latent vectors.

To test this, observers saw triplets of images corresponding to three vectors of equal length *r* in the GAN’s latent space (see Figure 3a). The observers’ task was to decide which one of two flanking stimuli was more similar to a centrally presented probe stimulus (see Figure 3b). Importantly, one of the two flanker stimuli (referred to as the “standard”) was orthogonal to the probe in latent space, while the other stimulus (the “target”) was rotated away from the standard towards the probe by a latent space angle *ϕ* (see Figure 3a). Thus, the target was closer to the probe in latent space.

**Figure 3:**
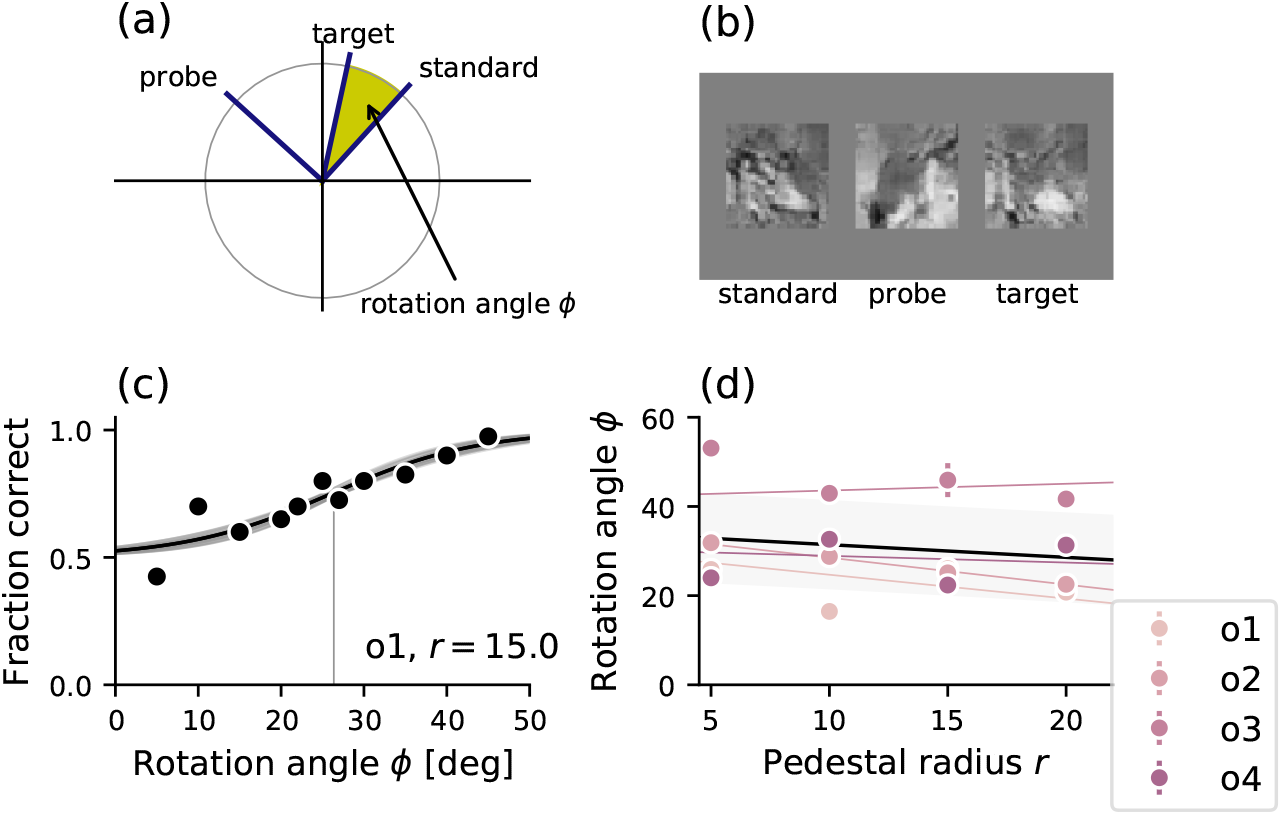
Sensitivity to latent space rotations is approximately constant. (a) On every trial, the observer sees three images that correspond to three latent space vectors of equal length *r*. While the standard and the probe vectors are orthogonal, the target vector is rotated from the standard towards the probe by an angle *ϕ* in the plane that contains the standard and probe vectors. Note that standard and probe vector pairs were chosen randomly on each trial, while the target was always constructed in the same way. (b) Example experimental display. Latent space vectors were projected to image space and observers had to choose the flanker that was more similar to the central probe image. The text below the gray display was not shown during the experiment. (c) Example psychometric function for discrimination of images created by latent space rotations for vectors of length *r* = 15. Each block represents 20 trials, the black line is the best fitting psychometric function, the thin gray lines correspond to samples from the posterior. The vertical gray line marks the estimated threshold for this observer. (d) Estimated rotation thresholds as a function of pedestal radius *r*. Each dot corresponds to a single threshold (error bars ± posterior standard deviation). The solid black line is the grand average across observers, the gray area is a 95% prediction interval for the grand average. Lightly colored lines are best fitting lines for different observers (see legend).

At a given length *r*, varying the latent space rotation *ϕ* traced out a psychometric function. For larger rotation angles, observers became better at detecting the target flanker, with performance reaching 75% typically between 20 and 30 degrees for observers o1, o2, and o4 (see Figure 3c for an example). Observer o3 had no prior experience with psychophysical experimentation and required larger rotations (45.91±2.21 degrees, mean ± sem).

Figure 3d shows 75% performance thresholds for all observers. The grand average straight line fit for the relationship between threshold and pedestal radius *r* had a slope of (−0.274 ± 2.676) and was not significantly different from 0 (Wald-test *z* = −0.106, n.s.).

### Homogeneity of tuning

The results in Figure 3d suggest that tuning for the multidimensional image manipulations which correspond to rotations in a GAN’s latent space might be invariant across the entire latent space. Unfortunately, measuring rotation discrimination thresholds throughout the 128 dimensions is experimentally not feasible^1^. We reasoned that if human tuning for latent space rotations changed across the GAN’s latent space, then being informed with the exact latent space locations of probe, standard and target should allow us to predict an observer’s trial-by-trial responses more accurately than being informed with just the angle of rotation. To test this idea, we trained a classifier to predict an observer’s single-trial responses from either (i) the full latent space representation of a trial, including the latent space rotations as well as the exact latent space locations of probe, standard and target for each single trial or (ii) a homogeneous feature set containing just the latent space rotations.

When predicting human responses from the homogeneous feature set, model predictions were marginally more consistent with human responses (average model-human consistency was 0.695±0.027, mean ± sem across 5 cross-validation folds, see Table 2) than they were when predicting human responses from the full feature set (0.672±0.016). For three out of our four observers, cross-validated consistency between human and classifier responses was larger if the classifier used the homogeneous feature set, although the advantage was typically small. For o3, the full features allowed for slightly better prediction of human responses but the difference was small and not significant (*t*(4) = 0.22, n.s.). Thus, in general, human responses in the second experiment were consistent with homogeneous tuning for latent space direction throughout the entire latent space learned by a GAN.

**Table 2:**
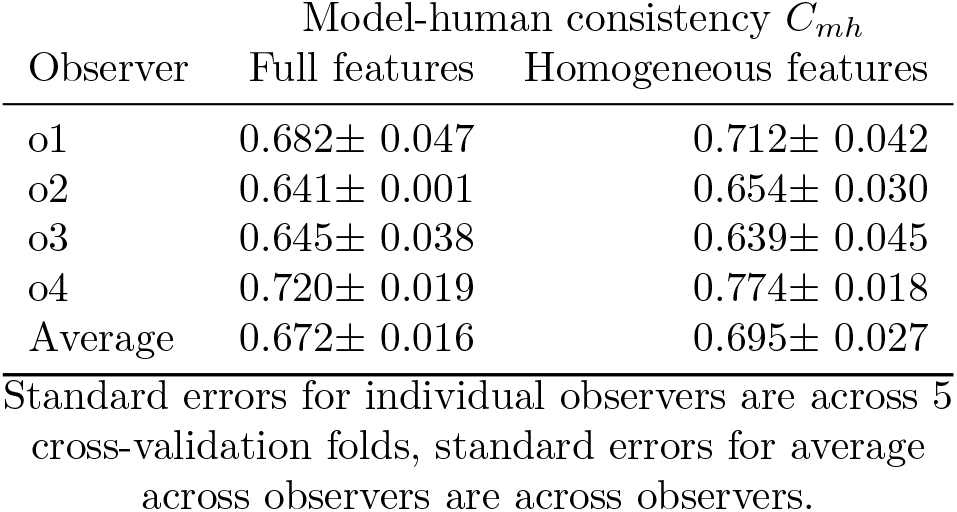
Homogeneity of trial-by-trial predictions across latent space locations.

### Independent sensitivity for radial and angular components predicts new discrimination data

We found that discrimination performance of images generated by a GAN can be described along two polar dimensions. Discrimination thresholds for images that correspond to different latent space radii follow a power law, while discrimination thresholds for images that correspond to different rotations of latent space vectors are approximately constant. Yet, it is unclear if these two independent factors can also describe discrimination performance if stimuli differ in both the radial and the angular components. Furthermore, we would like to know if these findings generalize to other experiments.

To address this question, we re-analyzed data from an independent, previously published study [20]. In that study, 8 observers decided which one of two flanking stimuli was closer to a centrally presented comparison. Crucially, one of the two flanking stimuli (the target) was constructed by adding noise to the latent space representation of the comparison, while the other was just another sample from the GAN. In that study, trial-by-trial responses were best predicted by comparing the Euclidean distance between the latent space representations of the target and the comparison to the Euclidean distance between the latent space representations of the non-target flanker and the comparison. We performed the same analysis here, but instead of the Euclidean distance, we used the sum of the difference in log radius weighted by the inverse radius discrimination threshold and difference in rotation weighted by the inverse rotation discrimination threshold. Note that some of the observers involved in the two studies differed, so we based this prediction on average radius and rotation discrimination thresholds as predicted by the respective mixture models.

In six out of eight observers, the threshold-weighted distance predicted trial-by-trial responses better than Euclidean distance. On average, threshold-weighted distance predicted trial-by-trial responses slightly better (area under the receiver operating curve: 83.7±0.9%, mean ± standard error) than for Euclidean distances (82.5±1.1%, paired T-test *t*(7) = 2.87*, p* = 0.02). Importantly, this was achieved without any additional free parameters and based on averages across observers in the current study.

## Discussion

We find that human discrimination performance in high dimensional feature spaces can be described by two independent factors; a contrast factor obeying a power law and a content factor with approximately constant sensitivity. We find that an image model based on deep neural networks—a GAN—recovers these two factors from natural images and represents them as high dimensional polar coordinates.

We find that discrimination thresholds for radius increments increase with an average exponent of 0.52, similar to values reported for grating contrast ([4] reports an average of 0.58, [30] report 0.59 for 2cpd gratings and 0.68 for 8cpd gratings). This is qualitatively and quantitatively consistent with the idea that the images’ global scene contrast reflects the latent space radius. On the other hand, tuning for the rotational component of the GAN’s latent space is unaffected by latent space radius, similar to what had previously been reported of more low level stimulus dimensions such as spatial frequency and orientation [4, 3].

Factorial representations of images have been proposed—more or less explicitely—before [31, 32, 33]. These accounts have typically discussed such representations in the context of contrast gain control and divisive normalization [34, 35, 36], arguing that disentagling contrast and other image properties could be accomplised by a system that represents images in polar coordinates with the radius component representing image contrast [31]. Our results imply that these ideas might hold for considerably more holistic image properties as well.

In recent years, a large number of GAN flavours and architectures have been developed [19, 37, 38, 39, 40, 41, 42, 21, 22] and it is not clear to what extent our findings carry over to these alternative GAN types. One architecture that has been particularly influential is the DCGAN architecture [37]. DCGANs are deep convolutional, purely feedforward networks with a generator that maps from the latent representation to image space. We qualitatively assessed a number of variations of DCGAN generators [42, 39, 40, 41] and found similar representations. On the other hand, modern GAN variations tend to use more elaborate ways of including latent variables [21, 22] and it is not directly obvious how our results would carry over to the other architectures. Thus, our results are expected to be directly applicable to DCGAN architectures but more work is needed to establish similar connections for other GAN variations.

Many characterizations of visual performance have been obtained with highly simplistic stimuli, such as gratings [3, 4, 28, 36, 27] or simple line drawings [43, 44, 45]. Parametric measurements of behavioural performance in the high-dimensional and non-linear feature space that defines natural-appearing images is difficult [20]. For example, Sebastian et al [46] superimposed target gratings on natural images resulting in stimuli with artificial foreground and natural background—still a very unnatural combination. Humans seem to be highly sensitive to deviations from natural appearance [47] and it is unclear to what extent deviations from natural appearance bias performance in psychophysical tasks involving natural images. Here, we used GANs to coordinate high-dimensional image manipulations and constrain them to remain approximately natural. Such coordinated high-dimensional stimulation may be particularly useful for characterizing multi-dimensional stimulus representation in physiological experiments [23]. It is well known that neural responses to natural input can be quite different from responses to synthetic stimuli [48] and generative image models such as GANs can help constrain stimulation to remain approximately natural.

## Methods

### Training the GAN

We trained a Wasserstein GAN [24] with DCGAN architecture [37] using gradient penalty [25] on the 60 000 images of the CIFAR10 dataset [26]. In brief, the trained GAN consisted of two parts, a generator network *G* and a discriminator network *D* that were trained by alternating between 5 parameter updates for the discriminator maximizing the error function and one parameter update for the generator minimizing this error function.

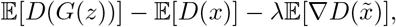

Here 𝔼[·] denotes expectation, *z* is a 128-dimensional isotropic standard normal variable, *x* are images from the training dataset, 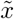 is an example image at a random location between the simulated images *G*(*z*) and the real images *x*, and ∇ denotes the gradient oper-ator (with respect to 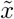). The details of the training procedure have been described elsewhere [20].

### Psychophysics

The same four observers gave informed consent to participate in both experiments. The experimental procedures were approved by the York University ethics board and were in accordance with the original declaration of Helsinki. Observers o1 and o2 were authors (ES and JP respectively), observers o3 and o4 were naive to the purpose of the experiment. Furthermore, observer o3 had no prior experience with psychophysical experimentation. Stimuli consisted of 32×32 pixel grayscale images presented on a linearized CRT monitor (Sony Triniton Multiscan G520) in an otherwise dark room. Maximum stimulus luminance was 106.9 cd/m^2^, minimum stimulus luminance was 1.39 cd/m^2^. The background luminance was set to medium gray (54.1 cd/m^2^). At a viewing distance of 56 cm, each generated stimulus image subtended approximately 1 degree visual angle. We re-rendered the stimuli on every frame using a random dithering method [49] to increase the luminance resolution of the 8-bit display. Stimuli were presented for 200 ms and were then removed from the screen. The observers then had 1000 ms to respond and they received feedback about their responses after every trial. If an observer didn’t respond within 1000 ms, the trial was marked invalid and excluded from analyses. Each measured psychometric function was based on 150–480 trials resulting in a total of 2560–3765 trials per observer across both experiments. We then fit the data for each psychometric function separately using a Weibull function in the radius increment detection experiment and a logistic function in the rotation discrimination experiment (both parametrized as in [50]). Psychometric functions were determined using Bayesian inference, which tends to give slightly better confidence intervals [51], by directly integrating the posterior [52]. Thresholds were defined as the point at which the psychometric function reached 75% correct performance.

### Linear mixed effects modeling

Observers’ radius discrimination thresholds were modelled using a linear mixed effects model of the form

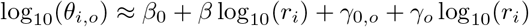

where *θ*_*i,o*_ is the estimated radius increment threshold for observer *o* at pedestal radius *r*_*i*_, and the *β* and *γ* are fixed and random effects parameters respectively. Thus, the average slope was given by *β* and the *γ*_*o*_ modelled the difference in slope between individual observer’s slopes and the average.

Rotation discrimination thresholds were similarly modelled using a linear mixed effects model of the form

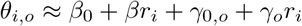

with in this case *θ*_*i,o*_ denotes the rotation threshold for observer *o* at pedestal radius *r*_*i*_.

### Trial-by-trial contrast analysis

To evaluate how much information other measures of contrast contributed to observers’ radius discrimination performance, we asked how well a linear classifier could predict the observers trial-by-trial choices. We trained logistic regression classifiers with ridge penalty to five different datasets and used 5-fold cross validation to adjust the strength of the ridge penalty. The first dataset contained just the radii presented on the left and right sides. The second dataset contained the Michelson contrast of the two presented images given by

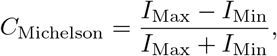

where *I*_Max_ and *I*_Min_ are the maximum and minimum pixel intensities of the presented stimulus. The third dataset contained the images’ RMS contrast given by the ratio between the standard deviation of a stimulus’ pixel values and the mean of a stimulus’ pixel intensities. The fourth dataset contained the images’ Weber contrast given by *C*_Weber_ = (*I*_Max_ − *I*_Min_)/*I*_Min_. Finally, the fifth dataset contained radii as well as all three contrast measures for both stimuli. Data from each observer were fit separately for this analysis.

### Trial-by-trial homogeneity analysis

To evaluate how much additional information was contained in the GAN’s full latent vectors, we used a classification analysis on two different sets of features. The *homogeneous* feature set consisted of the radius as well as sine and cosine of the rotation angles of the left and right flankers respectively for a total of 5 features. Note that on any trial, either the left or right rotation angle was 0. The *full* feature set consisted of the features from the homogeneous feature set as well as the full latent vectors of the left and right flankers and the probe. Thus, the full feature set had a total of 3×128 + 5 = 389 features. To be able to detect both latent space directions and regions of inhomogeneous sensitivity, we used a *C*-SVM with a radial basis function kernel [53] as implemented in scikit-learn [54] to predict the observers’ left or right choices. Cross-validated prediction accuracy was evaluated over 5 folds for a range of different kernel bandwidths and regularization strengths. For each model, we report the best cross-validated prediction accuracy over all combinations of kernel bandwidth and regularization strength.

## Data availability

Upon publication, all data and analysis code will be available upon at https://doi.org/10.5281/zenodo.3235639.

## Acknowledgements

This work was supported by the York University Bridging Fund.

## Footnotes

1 With a coarse sampling of 5 grid locations in each dimension, a 5-dimensional latent space would already require over one million discrimination thresholds to be estimated, and a 128-dimensional latent space would require ≈10^89^ discrimination thresholds!

